# Synergistic Electroceutical-Glucocorticoid Intervention Mitigates Dexamethasone-Induced Muscle Atrophy in Aging Skeletal Muscle

**DOI:** 10.64898/2026.03.10.709862

**Authors:** Min Young Kim, Sohae Yang, Jeongmin Kim, Yongkoo Lee, Minseok S. Kim

**Affiliations:** Department of New Biology, DGIST, Daegu 42988, Republic of Korea; Renal Division, Division of Engineering in Medicine, Department of Medicine, Brigham and Women’s Hospital, Harvard Medical School, Boston 02113, MA, USA; Department of Medical Device, Daegu Research Center for Medical Device and Green Energy, Korea Institute of Machinery & Materials (KIMM), Daegu 42994, Republic of Korea; Translational Responsive Medicine Center, DGIST, Daegu 42988, Republic of Korea; New Biology Research Center (NBRC), DGIST, Daegu 42988, Republic of Korea; CTCELLS Corp., Daegu 42988, Republic of Korea

**Author notes:** Corresponding author. E-mail address (M. S. Kim).

**Keywords:** aged skeletal muscle, dexamethasone, atrophy, electric medicine, electroceuticals

## Abstract

Dexamethasone (DEX), a synthetic glucocorticoid widely prescribed for allergic and inflammatory diseases, is known to induce adverse effects, particularly skeletal muscle atrophy. DEX-induced atrophy exacerbates sarcopenia and has a more pronounced impact on aged skeletal muscle than on young skeletal muscle. To address this unmet clinical need, we introduce an electroceutical approach that counteracts DEX-induced muscle atrophy and enhances functional recovery in aging muscle. When applied to both young and aged human-derived skeletal muscle cells (skMCs) exhibiting DEX-induced atrophy, electroceutical treatment promoted recovery of myotube diameter and upregulated hypertrophy-related gene expression. Furthermore, in a preclinical study, young and aged mice treated with DEX to induce muscle atrophy exhibited significant muscle recovery following electroceutical treatment. This effect was evident from the restored cross-sectional area (CSA) of type IIA muscle fibers and the upregulation of hypertrophy-related genes. This study highlights electroceuticals as a pioneering non-pharmacological strategy complementary to glucocorticoid therapies, potentially transforming clinical outcomes and quality of life, particularly for older populations vulnerable to muscle wasting.

## 1. Introduction

Glucocorticoids, particularly DEX, are widely prescribed for managing allergic and inflammatory diseases, especially among older adults ^1–3^. Although DEX effectively controls inflammation, it imposes a major clinical drawback by inducing skeletal muscle atrophy ^4^. This atrophic process is partly driven by increased protein degradation and suppressed protein synthesis via FoxO3 phosphorylation inhibition ^2, 5^. In older patients, these catabolic effects become more pronounced ^6^, exacerbating muscle wasting in populations already susceptible to sarcopenia ^7^. Consequently, the resulting deterioration in muscle mass not only compromises mobility but also reduces metabolic capacity and immune function, ultimately elevating mortality risk ^8–11^. Other synthetic glucocorticoids, such as prednisone, methylprednisolone, hydrocortisone, triamcinolone, and betamethasone, similarly induce muscle atrophy by comparable mechanisms ^12^. While clinicians attempt to balance therapeutic benefits against these adverse outcomes, no effective pharmacological therapy currently exists to fully prevent or reverse glucocorticoid-induced atrophy in older adults ^13^. Exercise is frequently recommended as a prophylactic measure ^8^, but it remains impractical for many older or hospitalized patients owing to limited physical capacity. Hence, there is a pressing need for non-pharmacological approaches that can be deployed safely in this vulnerable population ^14^.

The therapeutic landscape for sarcopenia remains barren, with zero FDA-approved pharmacological interventions despite decades of intensive research. Myostatin inhibition ^15^, initially considered the most promising approach, has encountered significant setbacks in clinical translation ^16–18^. Bimagrumab, a fully human monoclonal antibody targeting activin type II receptors, demonstrated muscle mass increases of 3.6% in Phase II trials but failed to achieve meaningful functional improvements in primary endpoints (6-minute walk distance, stair climbing power). More concerning, myostatin blockade strategies have been associated with serious adverse events including cutaneous malignancies (squamous cell carcinoma incidence: 4.2% vs 0.8% placebo), telangiectasia (18.7% incidence), and paradoxical muscle weakness in 12% of treated patients ^19–22^. Similarly, AMPK agonists and selective androgen receptor modulators have demonstrated limited efficacy with significant safety concerns, including hepatotoxicity and cardiovascular events ^23^. These failures underscore the critical need for alternative therapeutic modalities that can address muscle wasting without introducing additional pharmacological burden, particularly in older adults with polypharmacy concerns and increased drug interaction risks.

Considering these limitations, various non-pharmacological interventions have gained attention ^24–27^. Historically, electrical stimulation (ES) has been utilized in rehabilitation settings to augment muscle mass and strength ^28, 29^, and it can effectively induce muscle hypertrophy ^30–34^. Nevertheless, its therapeutic potential for drug-induced muscle atrophy, in aged skeletal muscle, remains underexplored. To address this gap, we developed high-throughput ES screening (HiTESS), a platform that enables systematic evaluation of electrical stimulation parameters on primary human skeletal muscle cells using only 5×10³ cells per condition ^27^.

Building on this foundation, we explored that specific ES conditions can recover the atrophy due to Dex treatment in aged skMC and we compared the restorative effect with young skMCs. The present study represents the first dedicated investigation of a combined electroceutical-pharmacological approach to mitigate DEX-induced muscle atrophy. As a result, we first evaluated whether electroceutical regimens could reverse DEX-induced atrophy in human-derived skMCs, quantified molecular recovery by profiling myogenesis-related gene expression using quantitative PCR, and subsequently translated these findings into an in vivo mouse model through computational simulation (CT-based 3D reconstruction) to optimize ES conditions for the tibialis anterior (TA) muscle (Fig. 1 and Fig. S1). Our results reveal that although DEX-induced muscle wasting is more severe in aged tissue, properly tuned electroceutical treatment substantially restores muscle fiber cross-sectional area, weight as well as volume. By highlighting the clinical feasibility of coupling electroceuticals with glucocorticoid therapy, our findings open a new avenue for non-pharmacological interventions designed to reduce drug-induced muscle atrophy, especially in older patients who cannot rely solely on exercise. We anticipate that this novel synergistic strategy will reshape future therapeutic protocols and encourage further research into the clinical translation of electroceutical technology.

**Fig. 1.**
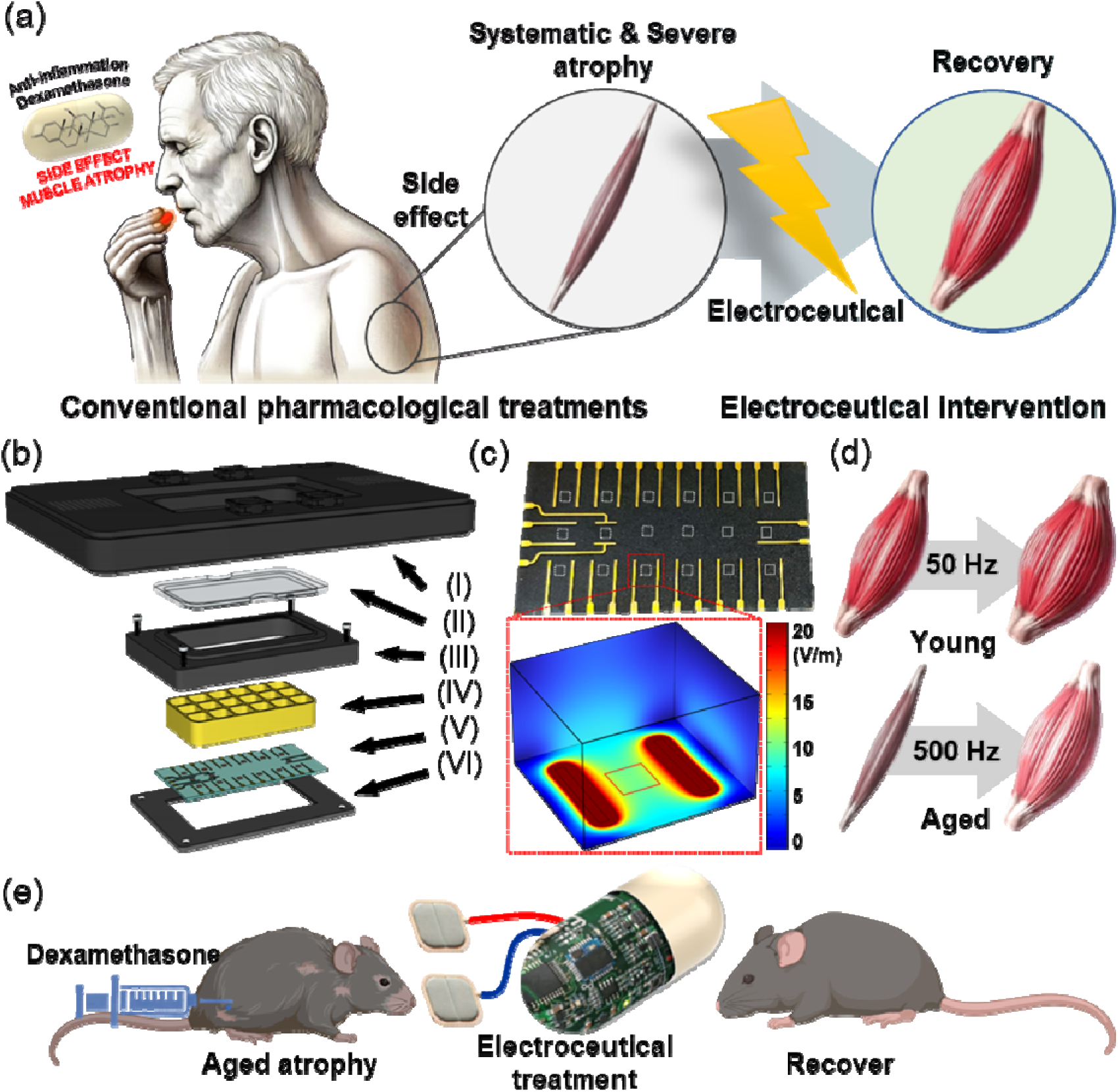
Age-tailored electroceutical platform reverses glucocorticoid-induced muscle wasting through frequency-optimized stimulation. (a) Muscles in aged individuals undergo significant atrophy because of DEX, yet the associated side effects may be attenuated through the synergistic application of electroceutical treatment. (b) Configuration of the HiTESS platform. (I) HiTESS pulse generator, (II) cover, (III) upper holder with connecting PCB, (IV) chamber with O-ring, (V) HiTESS substrate, and (VI) Bottom holder. (c) The HiTESS substrates with electric field simulation. (d) The effective ES frequency to recover the skMC was 50 Hz for young and 500 Hz for aged. (e) Dex-induced atrophy aged mouse model validated the efficacy of the electroceutical.

## 2. Results and Discussion

### 2.1. Differential Vulnerability of Young and Aged Skeletal Muscle to Dexamethasone-Induced Atrophy

To investigate the differential effects of DEX on muscle atrophy between young and aged skeletal muscle cells, we first characterized the senescent phenotype of our cell models and then examined their response to glucocorticoid-induced stress. During cellular senescence, many senescence-related molecules such as lipofuscin, senescence-associated β-galactosidase (SA-β-Gal), p16 and p53 are accumulated in aged cells ^35^. To confirm that the skMCs used in this study exhibited age-related characteristics, we performed a SA-β-Gal assay. The proportion of SA-β-Gal–positive cells was significantly higher in aged skMCs compared to young skMCs (Fig. S2), corroborating their senescent phenotype. This finding is consistent with previous reports demonstrating increased SA-β-Gal activity as a hallmark of cellular senescence in muscle tissue ^36^. The accumulation of senescent cells in aged muscle has been linked to reduced satellite cell function and impaired regenerative capacity, which may contribute to the increased susceptibility to atrophic stimuli observed in sarcopenia ^37^.

We evaluated how DEX at various concentrations influences both young and aged skeletal muscle cells by measuring myotube diameter and fusion index, key indicators of muscle differentiation. Immunostaining for myosin heavy chain 1 (Myh1) and nuclei [Fig. 2(a)] revealed a dose-dependent increase in atrophy for both age groups, though the aged control myotubes already exhibited a 38.82% smaller diameter compared to the young control. This baseline difference likely reflects the cumulative effects of aging on muscle fiber size and protein synthesis capacity. In the young group, 25, 50, and 100 μM of DEX reduced myotube diameter by 20.7%, 30.3%, and 38.2%, respectively, compared to the young control [Fig. 2(b)]. Aged skMCs were more severely affected, showing reductions of 29.1%, 38.2%, and 45.6% relative to the aged control.

**Fig. 2.**
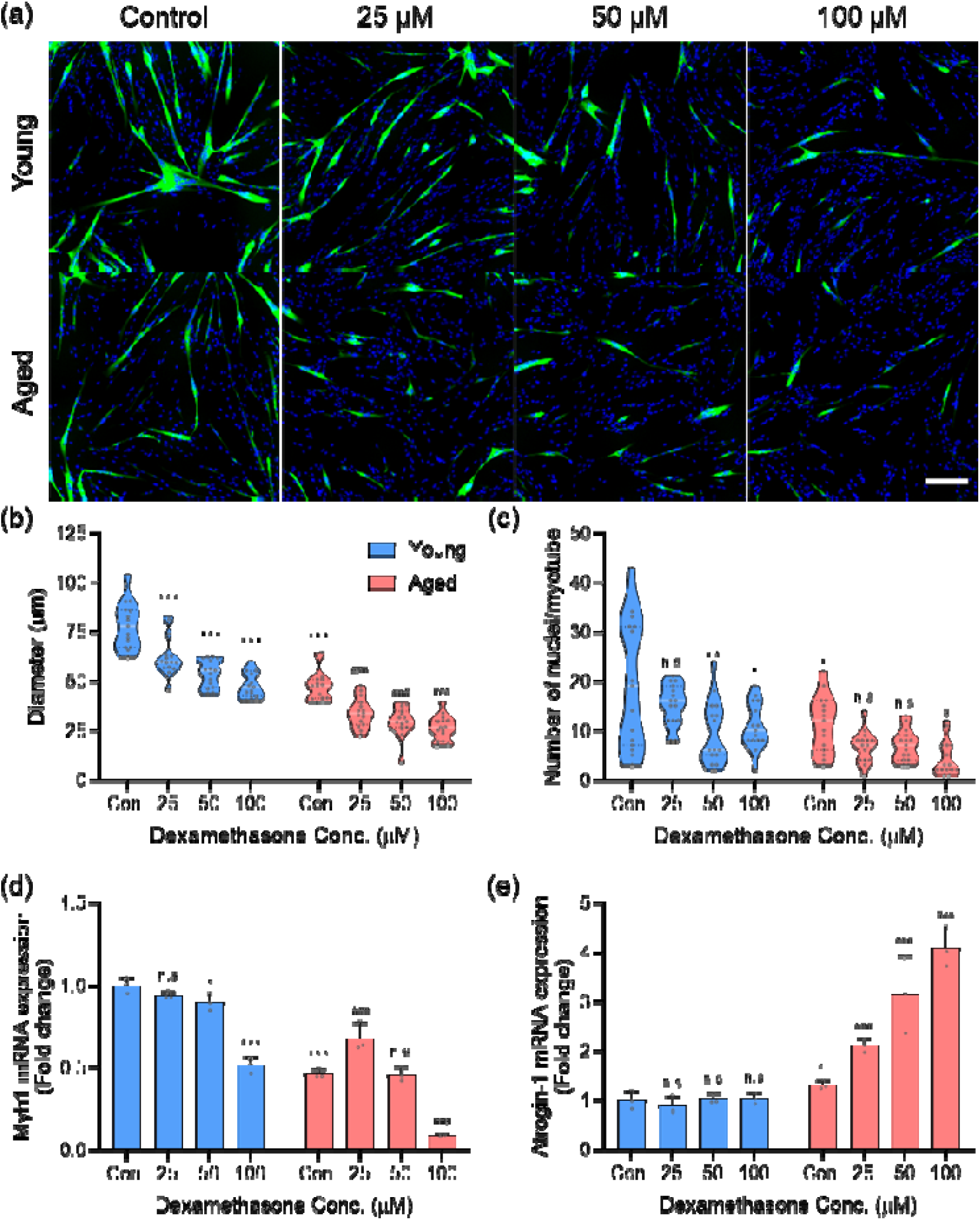
Comparison of the degree of atrophy in young and aged muscles as a function of DEX concentration. (a) Representative images of myotubes stained for Myh1 (green) and nuclei (blue) (scale bar: 200 μm). (b) Myotube diameter in young and aged myotubes following DEX-induced atrophy. (c) Fusion index in DEX-induced atrophy myotubes. Young myotubes exhibit a dose-dependent response, whereas aged cells show a dramatic decline in diameter and fusion index even at low DEX concentrations. (d, e) mRNA expression levels of Myh1 (d) and atrogin-1 (e) (n=3). (*\* P < 0.05; ** P < 0.01; *** P < 0.001 versus young control; # P < 0.05; ## P < 0.01; ### P < 0.001 versus aged control*)

The fusion index followed a similar trend [Fig. 2(c)], with aged myotubes exhibiting a 41.5% lower baseline than young controls and showing disproportionately higher declines under DEX treatment. The reduced fusion capacity in aged muscle cells may be attributed to altered expression of myogenic regulatory factors and fusion-related proteins, which are known to decline with age ^38^. This impaired fusion not only limits muscle regeneration but also compromises the muscle’s ability to maintain mass under catabolic conditions.

In line with these morphological data, Myh1 mRNA expression diminished significantly in both groups, with aged skMCs showing more pronounced reductions [Fig. 2(d)]. The suppression of Myh1, a major contractile protein, directly contributes to the loss of muscle mass and force-generating capacity. Conversely, atrogin-1, a muscle-specific E3 ubiquitin ligase and central mediator of protein degradation, was markedly upregulated [Fig. 2(e)]. The enhanced expression of atrogin-1 in aged muscle cells suggests an exaggerated activation of the ubiquitin-proteasome system, which has been implicated in age-related muscle wasting ^39^.

The mechanistic basis for this heightened vulnerability in aged muscle likely involves multiple pathways. Glucocorticoids are known to activate FoxO transcription factors, which induce atrogin-1 and MuRF1 expression ^40^. In aged muscle, this pathway may be primed for hyperactivation due to reduced IGF-1/Akt signaling, increased oxidative stress, and chronic low-grade inflammation ^41^. Additionally, aged muscle exhibits impaired mitochondrial function and reduced autophagy flux, which may compromise the cellular stress response to glucocorticoids ^42^.

Taken together, these data show that DEX triggers a disproportionately severe atrophic response in aged senescent muscle, characterized by reduced myoblast fusion, decreased MyH1 synthesis, and heightened proteasomal degradation. This age-related vulnerability to glucocorticoid-induced atrophy recapitulates clinical observations in elderly patients receiving glucocorticoid therapy, who often experience more rapid and severe muscle wasting compared to younger individuals. Understanding these age-specific responses is crucial for developing targeted interventions. The heightened susceptibility of aged muscle to DEX-induced atrophy provides a relevant model for evaluating potential therapeutic interventions, including the electroceutical approaches investigation.

### 2.2. Electroceutical Intervention Reverses Dexamethasone-Induced Muscle Atrophy Through Enhanced Myogenesis and Suppressed Proteolysis

Having established the heightened vulnerability of aged muscle to DEX-induced atrophy, we investigated whether ES, delivered via an electroceutical platform, could counteract these detrimental effects in both young and aged skeletal muscle cells. Our approach utilized a standardized ES protocol to evaluate its therapeutic potential across different age groups and DEX concentrations. As shown in Fig. 3(a), ES treatment resulted in marked improvements in myotube morphology compared to DEX-treated controls without ES [Fig. 2(a)]. The restorative effects were evident in both age groups but were particularly pronounced in aged muscle cells. Quantitative analysis of myotube diameter revealed that ES increased baseline myotube size by 2.3% in young controls (from 73.46±24.20μm to 75.11±23.99μm) and by 5.8% in aged controls (from 53.39±20.90μm to 56.48±19.16μm) [Fig. 3(b)]. This baseline enhancement suggests that ES promotes myogenic processes even in the absence of catabolic stress. When examining the therapeutic effects of ES under DEX treatment, we observed dose-dependent restoration of myotube diameter. In young muscle cells treated with 25 and 50 μM DEX, ES increased myotube diameter by 4.1% and 15.0%, respectively, compared to their DEX-only counterparts (25μM: 60.65μm vs 63.15μm; 50μM: 53.26μm vs 61.26μm). However, at 100 μM DEX, ES treatment resulted in a 7.9% decrease in myotube diameter (47.22μm vs 43.50μm). This observation in young cells under high-dose glucocorticoid exposure suggests that the combination of severe chemical stress and ES might have surpassed a tolerable physiological threshold. It is conceivable that at such high stress levels, the addition of ES, rather than promoting recovery, could have contributed to an excessive stress load, potentially leading to increased cellular vulnerability or even cytotoxicity. This highlights the possibility of a ‘therapeutic window’ for ES application, where its beneficial effects are prominent under moderate stress, but caution may be needed under conditions of extreme cellular duress. The aged muscle cells showed even more dramatic improvements, with ES increasing myotube diameter by 54.2% (33.15μm vs 51.13μm), 52.6% (28.90μm vs 44.09μm), and 66.8% (25.42μm vs 42.41μm) under 25, 50, and 100 μM DEX concentrations, respectively. Notably, ES treatment of aged cells exposed to 100 μM DEX restored myotube diameter (to 42.41±9.90μm) to levels approaching untreated aged controls (ES experiment control: 53.39±20.90μm), demonstrating strong therapeutic efficacy.

**Fig. 3.**
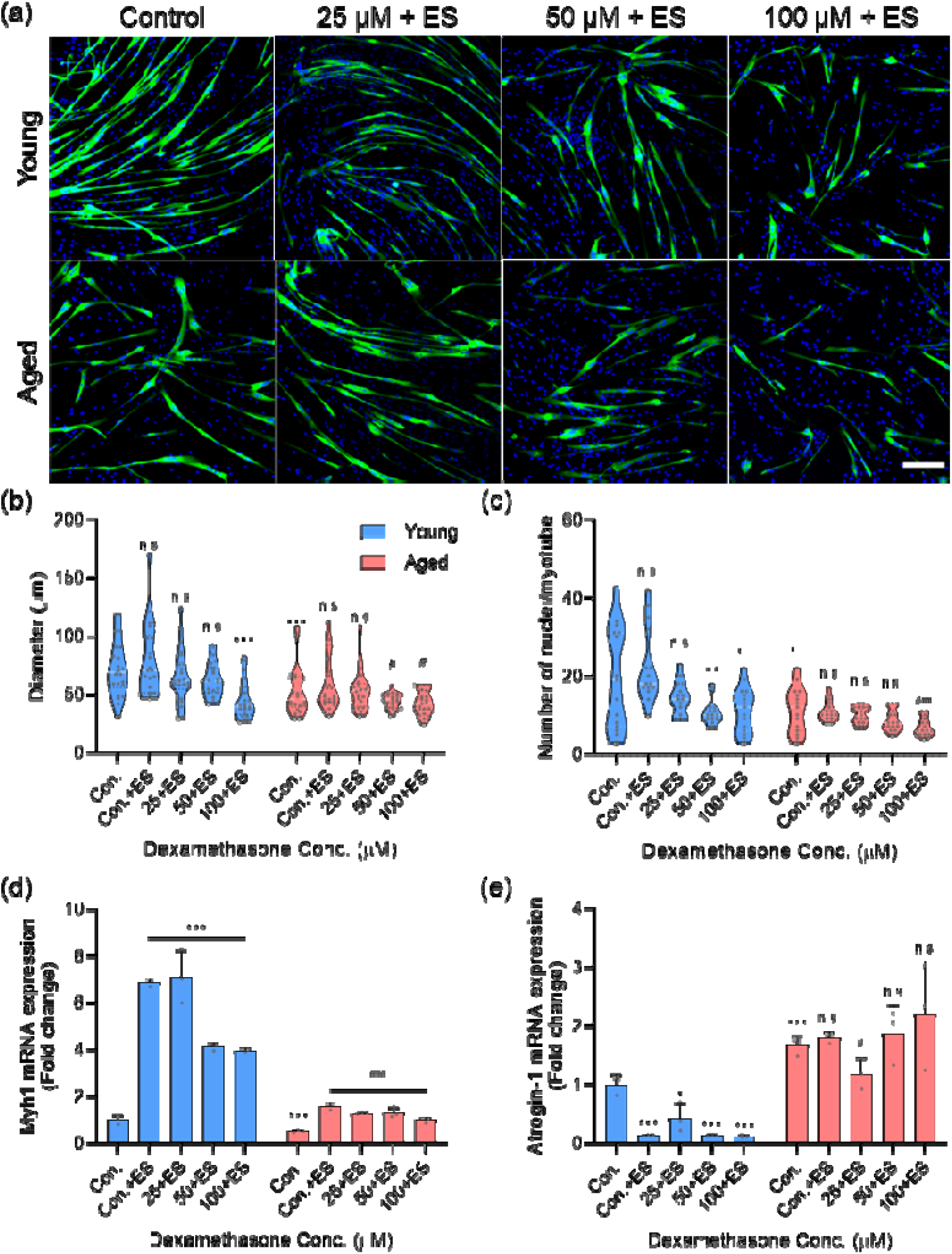
DEX-induced atrophy can be prevented by electroceutical treatment. (a, b) Representative images (a) and quantification (b) of myotubes stained for Myh1 (green) and nuclei (blue) (scale bar: 200 μm). (c) Fusion index in DEX-induced atrophy myotubes. (d, e) mRNA expression levels of Myh1 (d) and atrogin-1 (e) (n=3). (*\* P < 0.05; ** P < 0.01; *** P < 0.001 versus young control; # P < 0.05; ## P < 0.01; ### P < 0.001 versus aged control*)

The fusion index data revealed differential age-specific responses to ES treatment [Fig. 3(c)]. In young muscle cells, ES produced modest improvements in fusion capacity, increasing the fusion index from 20.50±13.08% to 23.43±9.23% in controls. Under DEX treatment, the effects of ES on fusion indices in young cells varied with DEX concentration; a slight increase was observed at 100 μM DEX, while slight decreases were noted at 25 μM and 50 μM DEX. However, the findings in aged muscle cells regarding ES effects on fusion capacity differed from initial expectations. In aged control cells, ES treatment resulted in a slight decrease in the fusion index from 12.00±5.42% to 10.71±2.13%, representing a 10.8% decrease [Fig. 3(c)]. Furthermore, under high-dose DEX (100 μM) treatment with ES, the fusion index for aged cells was 6.86±2.07%. This value was not maintained above the baseline aged control level of 12.00±5.42%. These particular data points suggest that further investigation is needed to fully understand the impact of ES on age-related deficits in myoblast fusion under these specific conditions.

To elucidate the molecular mechanisms underlying these morphological improvements, we analyzed the expression of key myogenic and atrophic markers. Myh1 mRNA expression analysis revealed that ES significantly upregulated this essential contractile protein in both age groups [Fig. 3(d)]. In young muscle cells, ES increased Myh1 expression by 6.2-fold in controls and maintained higher expression levels across all DEX concentrations compared to non-stimulated counterparts. The effect was even more pronounced in aged muscle cells, where ES induced a 3.2-fold increase in baseline Myh1 expression and sustained significantly higher levels even under 100 μM DEX treatment. This enhanced expression of structural proteins directly correlates with the observed increases in myotube diameter.

Conversely, analysis of atrogin-1 expression demonstrated that ES effectively suppressed proteolytic pathways activated by DEX [Fig. 3(e)]. In young muscle cells, ES reduced atrogin-1 expression by 54.0% (25 μM DEX), 62.6% (50 μM DEX), and 75.3% (100 μM DEX) compared to DEX-only counterparts. The suppressive effect was also observed in aged muscle cells, where ES decreased atrogin-1 levels by 37.4% in the 100 μM DEX group compared to DEX-only counterparts. This dual action, promoting anabolic processes while inhibiting catabolic pathways, reflects a broad therapeutic mechanism. The enhanced responsiveness of aged muscle to ES treatment warrants further discussion. Several mechanisms may contribute to this age-specific benefit. First, aged muscle exhibits reduced baseline electrical excitability due to alterations in membrane potential and ion channel expression. Exogenous ES may compensate for these deficits, restoring calcium transients necessary for proper muscle function. Second, ES has been shown to activate satellite cells and promote their fusion with existing myofibers, a process that is particularly impaired in aged muscle. The improvement in fusion index observed in DEX-treated aged cells with ES (compared to DEX-only treatment) might support this mechanism. Furthermore, ES may counteract age-related metabolic dysfunction. Electrical stimulation enhances mitochondrial biogenesis through PGC-1α activation, potentially offsetting the mitochondrial impairment characteristic of aged muscle. Additionally, ES can modulate inflammatory signaling pathways, reducing the chronic low-grade inflammation that primes aged muscle for enhanced glucocorticoid sensitivity. The clinical implications of these findings are substantial. Glucocorticoid-induced myopathy represents a significant clinical challenge, particularly in elderly patients who require chronic corticosteroid therapy for inflammatory conditions. Our data suggest that electroceutical interventions could provide a non-pharmacological approach to prevent or treat steroid-induced muscle wasting. The greater therapeutic benefit observed in aged muscle is particularly encouraging, as this population bears the highest burden of both sarcopenia and glucocorticoid-related complications.

Taken together, these data show that electroceutical treatment counteracts DEX-induced muscle atrophy through combined effects on myotube morphology, fusion capacity, and molecular signaling. The enhanced therapeutic response in aged muscle cells, which exhibit the greatest vulnerability to glucocorticoid-induced damage, points to ES as a targeted intervention for age-related muscle wasting. These findings support further investigation of electroceutical approaches in clinical settings, particularly for older patients at risk for iatrogenic muscle loss.

### 2.3. Electroceuticals Mitigate DEX-Induced Atrophy in Young and Aged Mice

The in vivo results revealed that electroceutical technology effectively mitigates DEX-induced muscle atrophy, with a particular focus on aged skeletal muscle. In vivo experiments were conducted using young and aged mice to evaluate this efficacy [Fig. 4]. DEX was administered to induce muscle atrophy, and subsequently, the mice were treated with electroceutical stimulation daily for 30 minutes over a period of six weeks. The TA muscle was the primary focus, with muscle CSA, mass, and volume being the key metrics evaluated. To verify muscle volume changes through objective imaging modalities, we calculated the volume of the TA muscle through CT scans [Figs. 5(a, b)]. The DEX group showed a volume reduction of 12.0% and 21.8% in young and aged mice, respectively, demonstrating age-dependent differential susceptibility to glucocorticoid-induced atrophy. However, with ES treatment, the young group exhibited an 11.3% increase in TA muscle volume compared to the DEX group, reaching levels like the control group, while the aged group demonstrated an 18.4% recovery compared to the DEX group, with only a 3.4% reduction in volume relative to the aged control group.

**Fig. 4.**
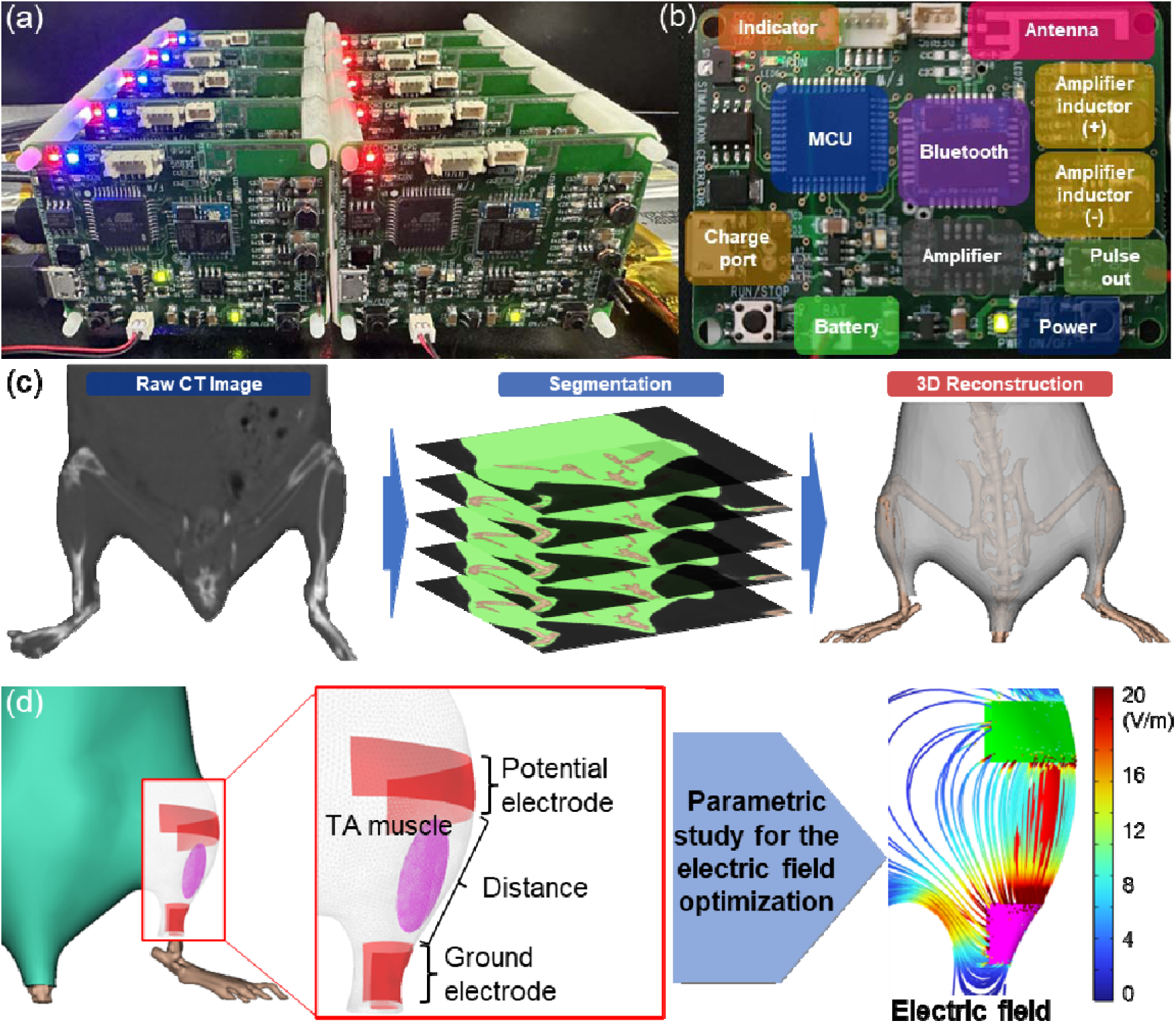
Electroceutical prototype and simulation for translating in vitro conditions to in vivo applications. (a, b) Representative images of the electroceutical device showing the electronic component layout. (c) 3D reconstruction of mouse hindlimb from micro-CT data showing TA muscle, gastrocnemius muscles, and bone structures for simulation. (d) Multiphysics simulation of E-field distribution in mouse hindlimb. Optimal electrode configuration (2-mm stimulating, 3-mm ground, 8-mm spacing) achieved target E-field of 11.8 V/m in TA muscle, matching in vitro conditions with 99.2% similarity. Color map indicates E-field intensity gradients.

**Fig. 5.**
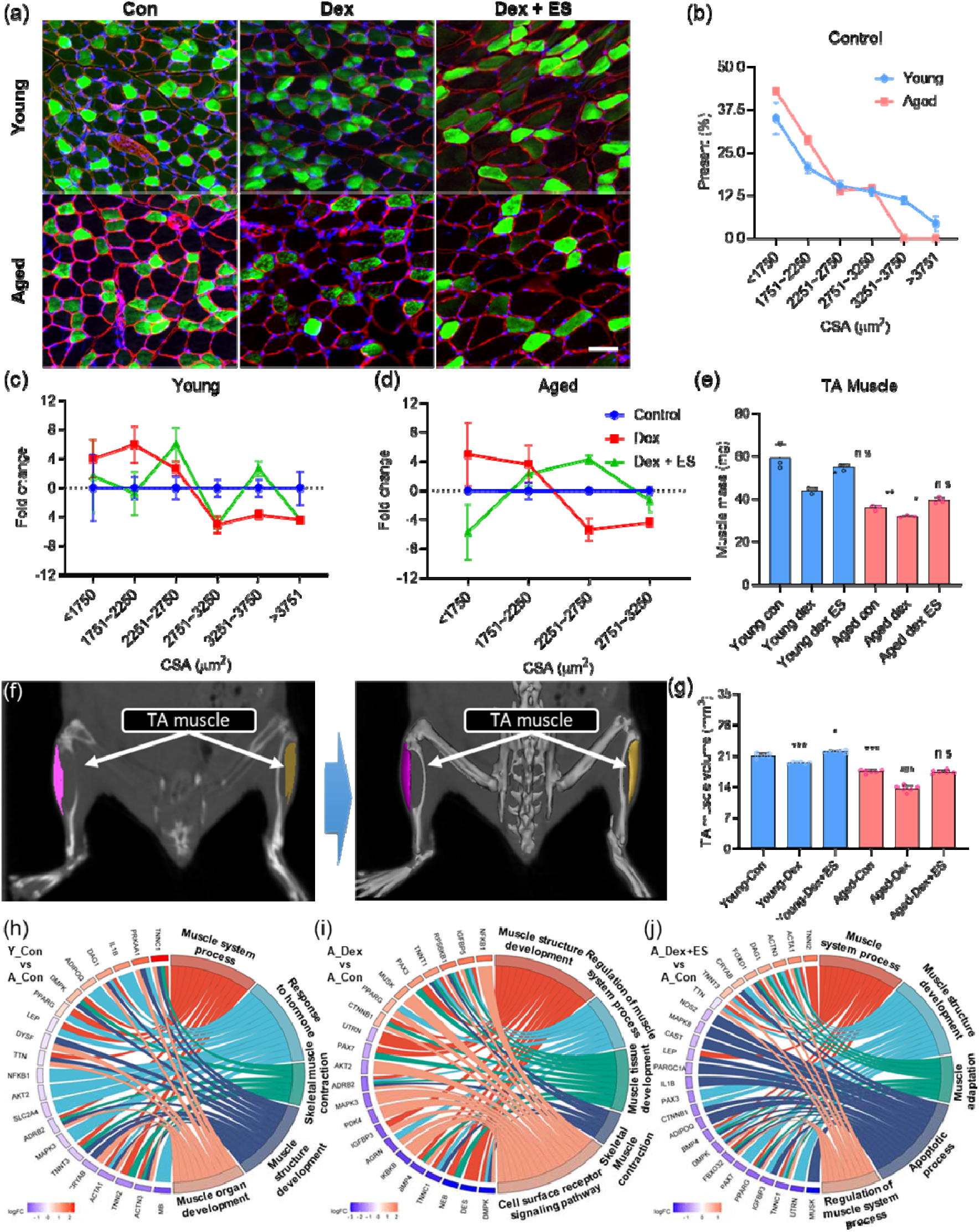
Validation of electroceutical treatment in young and aged mice. (a, b) TA muscle volume measurements from CT images (a) and corresponding quantification (b). (c) Quantification of TA muscle weight. (d) Representative images of CSA of Type IIA muscle fibers in young and aged mice (Blue: Nuclei, Green: Type IIA fibers, Red: laminin) (scale bar: 200 μm). (e-g) Quantification of myofiber CSA (e) comparison between young and aged controls, (f) comparison among young control, DEX-treated, and DEX+ES groups; (g) comparison among aged control, DEX-treated, and DEX+ES groups (n=3). (h-j) Myogenesis-related qPCR array results: (h) young control versus aged control; (i) aged DEX-treated group versus aged control; and (j) aged DEX with electroceutical-treated group versus aged control. (*\* P < 0.05; ** P < 0.01; *** P < 0.001 versus young control; # P < 0.05; ## P < 0.01; ### P < 0.001 versus aged control*)

Subsequently, we measured the mass of the TA muscle to corroborate the volumetric findings [Fig. 5(c)]. In both the young and aged groups, the DEX-treated group showed a reduction in TA muscle mass of 25.9% and 12.0%, respectively. Notably, the magnitude of mass reduction differed from volumetric measurements, with young mice exhibiting greater mass loss despite smaller volume reduction, potentially indicating age-related differences in muscle fiber density or composition. However, in the DEX+ES group, the young group showed an 18.6% increase in mass compared to the DEX group, confirming that atrophy was significantly alleviated through ES treatment. In the aged group, a 20.4% recovery was observed in the DEX+ES group compared to the DEX group, resulting in TA muscle mass that exceeded those of the control group, suggesting that ES provides not merely restoration but enhancement of muscle mass beyond baseline levels.

The muscle fiber CSA analysis provided critical insights into the cellular mechanisms underlying these macroscopic changes [Fig. 5(d)]. Representative images of Type IIA muscle fibers (Blue: Nuclei, Green: Type IIA fibers, Red: laminin) demonstrated clear morphological alterations across treatment conditions. The muscle fiber CSA showed a significant reduction, with aged mice experiencing a greater decline in larger muscle fibers (CSA > 2751 µm²) compared to young mice [Figs. 5(e-g)]. Specifically, aged mice treated with DEX exhibited a significant reduction in myotube diameter over CSA > 2251 µm². However, ES considerably counteracted these effects, leading to an increase in muscle fiber size between 2251 to 2750 µm² in aged mice. In the young group, DEX treatment resulted in a 4% reduction in muscle fiber area for fibers with CSA above 2751 µm². However, ES treatment increased the proportion of fibers in the 2251-2750 µm² and 3251-3750 µm² range. This confirms that ES treatment can alleviate the atrophy caused by DEX through preservation and restoration of myofiber cross-sectional dimensions. Therefore, our in vivo experiments demonstrate that the combination of pharmacological and electroceutical therapy can offset the atrophy induced by the side effects of DEX, highlighting the potential of this combined approach.

### 2.4. Gene Ontology Alterations in Skeletal Muscle in Response to DEX and ES

Gene Ontology analysis revealed profound age-related alterations in skeletal muscle gene expression patterns, providing molecular insights into the differential responses to DEX treatment and ES intervention. Comparative analysis between young and aged controls revealed significant shifts in gene expression profiles [Fig. 5(h)]. The upregulation of IL1B, a pro-inflammatory cytokine that serves as a key mediator of age-related chronic inflammation (inflammaging), indicated a heightened inflammatory state characteristic of sarcopenia. Concurrently, the upregulation of PPARG, a master regulator of lipid metabolism and adipogenesis, suggested metabolic reprogramming toward lipid accumulation, which is consistent with age-related intramuscular fat infiltration observed in sarcopenic muscle. These findings align with recent transcriptomic studies demonstrating that aging skeletal muscle exhibits a pro-inflammatory and metabolically dysregulated phenotype. The downregulation of structural genes such as ACTA1 (α-skeletal actin) and ACTN3 (α-actinin-3), both critical components of the sarcomeric apparatus essential for muscle contraction, pointed to compromised contractile machinery in aged skeletal muscle. This molecular signature is particularly significant as ACTN3 deficiency has been associated with reduced muscle power and increased susceptibility to muscle damage. GO enrichment analysis corroborated these findings, showing significant downregulation in muscle system processes and skeletal muscle contraction pathways, highlighting the decline in muscle functionality with aging at the transcriptional level.

DEX treatment in aged skeletal muscle induced a comprehensive downregulation of genes essential for muscle integrity and function [Fig. 5(i)]. The suppression of ADRB2 (β2-adrenergic receptor), a key mediator of catecholamine-induced muscle hypertrophy and metabolic activation, represents a critical mechanism by which glucocorticoids impair muscle anabolic signaling. Similarly, the downregulation of BMP4 (bone morphogenetic protein 4), a member of the TGF-β superfamily involved in myoblast proliferation and differentiation, indicates compromised muscle regenerative capacity. The suppression of structural proteins including DES (desmin), an intermediate filament protein crucial for sarcomeric integrity, and DMPK (dystrophia myotonica protein kinase), involved in muscle fiber stability and ion channel regulation, further exacerbates muscle atrophy and functional decline. This extensive downregulation creates a molecular environment conducive to accelerated muscle wasting and metabolic dysregulation. Paradoxically, the upregulation of CTNNB1 (β-catenin), a central component of the Wnt signaling pathway involved in cell adhesion and muscle regeneration, and IGFBP5 (insulin-like growth factor binding protein 5), a modulator of IGF-1 bioavailability, reflected a complex compensatory stress response. These upregulations may represent an attempt to maintain cellular integrity and growth signaling under glucocorticoid stress. GO analysis revealed downregulation in muscle structure development and regulation of muscle system processes, while pathways involving cell surface receptor signaling were upregulated, indicating a stress response aimed at maintaining cellular signaling integrity despite compromised structural components.

ES treatment in aged skeletal muscle demonstrated remarkable therapeutic potential by inducing upregulation of structural and functional genes [Fig. 5(j)]. The restoration of ACTA1 and ACTN3 expression represents a direct reversal of age-related contractile protein deficits, potentially enhancing muscle integrity and force-generating capacity. This finding is particularly significant as these proteins are fundamental to the excitation-contraction coupling process and their restoration could translate to improved muscle performance. The concurrent downregulation of key inflammatory mediators, including IL1B and NOS2 (inducible nitric oxide synthase), demonstrated ES’s potent anti-inflammatory effects. NOS2 downregulation is particularly important as excessive nitric oxide production contributes to muscle wasting through protein nitrosylation and mitochondrial dysfunction. Interestingly, the downregulation of BMP4 and DMPK in the ES-treated group, while initially counterintuitive, may reflect a normalization of stress-induced compensatory responses, suggesting that ES treatment reduces the need for these stress-responsive pathways. GO analysis revealed upregulation in muscle system processes and apoptotic process pathways, indicating improved muscle function and potentially enhanced clearance of damaged cellular components through regulated cell death mechanisms.

## 3. Conclusion

We report an approach that integrates electroceutical treatment with a commonly prescribed anti-inflammatory drug, DEX, to combat skeletal muscle atrophy, especially in aging populations where muscle recovery is already compromised. Our in vitro results demonstrate that electroceutical regimens substantially mitigate DEX-induced atrophy by enhancing protein synthesis and suppressing proteolysis, with a particularly strong protective effect in aged muscle cells. Through 3D computational simulations, we translated these ES parameters into an in vivo mouse model and confirmed that daily electroceutical interventions prevent or even reverse muscle fiber atrophy in DEX-treated mice. Notably, the impact of ES was most pronounced in older animals, suggesting that this combined therapy holds special promise for older adults who often have limited exercise capacity. By showing that non-pharmacological, electroceutical treatment can offset the atrophic side effects of a potent glucocorticoid, our findings introduce a completely new direction for drug–device synergy in musculoskeletal medicine. This strategy offers a tangible alternative to prescribing additional medications solely to counteract DEX. Future work should focus on mechanistic studies to clarify the signaling pathways modulated by ES, along with clinical trials to evaluate the feasibility and long-term safety of this combined approach in diverse patient populations. afflicting aged individuals who are most susceptible to detrimental drug side effects and functional decline.

## 4. Experimental Section

### 4.1. Cell Culture, DEX Treatment, and Electroceutical Application on Human-Derived skMCs

Primary skMCs were obtained from male donors aged 17 and 68 years (SK-1111, Cook Myosite, PA). Cells were expanded in T75 flasks utilizing a myotonic basal medium (MB-2222, Cook Myosite, PA) with 10% myotonic growth supplement (MB-3333, Cook Myosite, PA) and 1% antibiotic-antimycotic solution (SV30079.01, Cytiva, MA). After two days of proliferation, cells were harvested using 0.05% trypsin-EDTA (T3924, Sigma, MO). Subsequently, cells were collected by centrifugation at 1200 rpm for 3 minutes, and 5.0E03 cells were seeded into each ES well. To initiate cell differentiation, the medium was substituted with a differentiation medium (MB-5555, Cook Myosite, PA). After an additional two days, the cells were exposed to various concentrations of DEX (D4902, Sigma, MO). Two days later, electroceutical treatment was applied using the HiTESS platform, consisting of 2 hours of ES per well. Myotubes were subjected to ES at frequencies of 50 and 500 Hz with a pulse width of 0.01 ms, based on prior findings that suggest different frequencies optimally stimulate young (50 Hz) and aged skeletal muscle (500 Hz) recovery ^27^. These specific parameters were selected to target distinct recovery capacities in young versus aged muscle cells.

### 4.2. Immunocytochemistry and Senescence-Associated **β**-Galactosidase (SA-**β**-Gal) Assays

Myotubes were fixed in 4% formaldehyde (28908, Invitrogen, CA) for 15 minutes, then permeabilized with 0.5% Triton™ X-100 (T9284, Sigma, MO). After washing Dulbecco’s Phosphate Buffered Saline (DPBS) (LB001-02, Welgene, S. Korea), the cells were blocked in 2% Bovine serum albumin (BSA) (A3311, Sigma, MO) to prevent nonspecific binding. After blocking, the samples were washed with 0.5% BSA and incubated with primary antibodies (1:200 MF-20, DSHB, IA) for 2 hours at room temperature (RT). Following this, the myotubes were washed three times with 0.5% BSA and incubated with secondary antibody (1:200 A-11001, Invitrogen, CA) for 1 hour at RT. After incubation, the samples were washed five times in DPBS and mounted with antifade mounting medium containing DAPI (H-1200, Vector Laboratories, CA). Fluorescence images were captured using a Ti2 fully automated confocal microscope (Nikon, Japan). Image acquisition was performed using NIS software, and parameters such as myotube diameter and the number of nuclei per myotube were quantified using IMARIS Software (Bitplane, Switzerland).

The SA-β-Gal staining was conducted in accordance with the manufacturer’s specifications utilizing an SA-β-Gal assay kit (9860, Cell Signaling Technology, MA). SA-β-Gal levels were positively correlated with cellular senescence, as confirmed by the immunostaining procedure. Imaging was performed using an inverted microscope (DMi1, Leica, Germany) for subsequent quantification, and the number of SA-β-Gal-positive skMCs was assessed using IMARIS software.

The isolated TA muscle was embedded in an Tissue-Tek^®^ optimal cutting temperature (OCT) compound (4583, Sakura Finetek, CA) for cryosectioning. The muscle was sectioned into 7 µm-thick serial sections using a cryomicrotome (CM3050S, Leica, Germany). For the analysis of fiber size, a sequential staining procedure was applied. The sections underwent fixation and permeabilization, followed by incubation with antibodies against laminin (1:200 PA1-16730, Abcam, UK) and myosin heavy chain Type IIA (1:200 SC-71, DSHB, IA) for 2 hours. The sections were then washed three times in DPBS for 3 minutes each and incubated in the dark with secondary antibodies (1:200 A-11001, A-11012, Invitrogen, CA) for 1 hour. Images were acquired using a Nikon Ti2 confocal microscope to evaluate the fiber CSA. The quantification of myofiber area was performed using ImageJ software.

### 4.3. Real-time PCR

The myotubes were lysed in TRIzol™ (15596026, Thermo Fisher, MA), and RNA isolation was performed using chloroform and isopropyl alcohol according to the manufacturer’s instructions. cDNA synthesis was performed using TOPscript™ RT DryMIX (RT200, Enzynomics) on a MiniAmp™ Thermal Cycler (Applied biosystems, CA). Quantitative PCR was conducted on a StepOne™ Real-Time PCR System (Applied biosystems, CA) using KAPA SYBR FAST qPCR Master Mix (KK4602, Kapa Biosystems, MA), according to the manufacturer’s instructions. Gene expression was quantified using β-actin as the endogenous control. Primer sequences are provided in Table S2.

### 4.4 Multiphysics Simulation for In Vitro to In Vivo Translation

To ensure accurate translation of our in vitro electrical stimulation parameters to the in vivo setting, we conducted comprehensive multiphysics simulations to optimize electrode configuration and placement. First, we acquired high-resolution micro-CT images of mouse hindlimb musculature using a Quantum GX μCT system (Perkin Elmer, CT). The acquired DICOM images were imported into Mimics Innovation Suite (Materialise, Belgium) for segmentation and 3D reconstruction. Individual anatomical structures, including the TA muscle, surrounding muscles, and bone tissues, were carefully segmented based on their distinct radiodensities to create accurate 3D surface meshes.

The reconstructed 3D model was then imported into COMSOL Multiphysics software (COMSOL Inc., MA) for finite element analysis of electric field distribution. We assigned tissue-specific electrical properties to each anatomical structure based on published values (Table S1), ^43^ including electrical conductivity and relative permittivity for muscle, bone, and surrounding tissues. The TA muscle model had dimensions of 1.5 mm width, 3.5 mm length, and 0.75 mm thickness, representing typical measurements for adult mouse TA muscle

Our simulation strategy focused on identifying electrode configurations that would generate an electric field intensity of approximately 11.8 V/m within the TA muscle volume, matching our validated in vitro conditions. We systematically evaluated multiple electrode parameters through parametric sweeps: electrode widths ranging from 1-3 mm for both stimulating potential and ground electrodes, and inter-electrode distances from 6-10 mm [Fig. S1]. The applied voltage was maintained constant at 3.7 V to match in vitro conditions.

The simulation results revealed clear relationships between electrode geometry and resulting field distributions. Increasing electrode surface area enhanced electric field strength and current density within the target muscle, while greater inter-electrode distances attenuated these parameters following expected electromagnetic principles. Through iterative optimization, we identified several electrode configurations capable of producing the target field strength. For example, a configuration using 1-mm electrodes at 7-mm spacing generated 12.0 V/m, while 2-mm potential and 3-mm ground electrodes at 9-mm spacing yielded 11.7 V/m within the TA muscle volume. Based on comprehensive analysis of field homogeneity, current density distribution, and practical considerations for in vivo application, we selected the optimal configuration: a 2-mm wide stimulating electrode paired with a 3-mm wide ground electrode, positioned 8 mm apart. This arrangement achieved an average electric field of 11.6 V/m within the TA muscle, representing 99.2% similarity to our target in vitro conditions. Additionally, this configuration provided uniform field distribution throughout the muscle volume while minimizing current density hotspots that could cause tissue damage.

These simulation results provided critical guidance for electrode fabrication and placement protocols in subsequent in vivo experiments, ensuring faithful translation of our validated in vitro electroceutical parameters to the more complex in vivo environment.

### 4.5 Animal Model and Experimental Procedures

Female C57BL/6J young (6-months old) and aged (24-months old) mice were used. The mice were purchased from the Korea Research Institute of Bioscience and Biotechnology Laboratory Animal Resource Center. These animals were kept in a vivarium under a controlled 12-hour light-dark cycle with unlimited access to water and food. Mice were randomly divided into three groups to receive daily intraperitoneal injections with 0.1 mL of saline (control) or Dex (10 mg/kg in saline) for one week. Anesthesia was induced and maintained with 2% isoflurane for induction and 1% for maintenance (VIP 3000, MIDMARK). Considering the recovery capabilities of the aged mice, the maximum duration for anesthesia was restricted to 40 minutes. Accordingly, ES was administered daily for 30 minutes over a period of six weeks. Prior to electrode attachment, hair was removed from the mice’s legs using a hair removal cream. To facilitate ES application, the mice were restrained on a plastic plate, and electrodes connected to a electroceutical device were affixed to their legs. All animal experiments were conducted in accordance with the guidelines of the Institutional Animal Care and Use Committee (IACUC) of DGIST (approval number: DGIST-IACUC-21021001-0001).

### 4.6 Statistical Analysis

All data are presented as mean ± standard deviation (SD). Statistical analyses were performed using GraphPad Prism 9.0 (GraphPad Software, CA, USA). Comparisons between multiple groups were analyzed using one-way ANOVA followed by Tukey’s post-hoc test for multiple comparisons. Statistical significance was set at p < 0.05. All experiments were performed in triplicate with at least three independent biological replicates.

## CRediT Authorship Contribution Statement

**Min Young Kim:** Writing – review & editing, Writing – original draft, Visualization, Conceptualization, Methodology, Investigation, Data curation. **Sohae Yang:** Writing – review & editing. **Jeongmin Kim:** Software. **Yongkoo Lee:** Software. **Minseok S. Kim:** Writing – review & editing, Writing – original draft, Supervision, Conceptualization, Project administration, Funding acquisition.

## Acknowledgements

This work was supported by the National Research Foundation (NRF) of Korea, funded by the Korean government (MSIP) (No. 2022R1A2C2091870).

## Appendix A. Supplementary data

Supplementary data to this article can be found online at Biomaterials DOI

## Data Availability

Data will be made available on request.

