## Supplementary material for "Synergistic Electroceutical-Glucocorticoid Intervention Mitigates Dexamethasone-Induced Muscle Atrophy in Aging Skeletal Muscle": SI

** Corresponding author.*

**Table S1. Material properties for simulation**

|  | **Density (kg/m3)** | **Dielectric const. Electric**  **conductivity (S/m)** | **Dielectric const. Electric**  **conductivity (S/m)** |
| --- | --- | --- | --- |
| **Skeletal muscle** | 1090 | 4.35E+05 | 4.05E-01 |
| **Skin** | 1109 | 1.14E+03 | 1.22E-03 |

**Table S2. List of primers used for RT-PCR**

|  | **Forward primer sequence (5'-3')** | **Reverse primer sequence (5'-3')** |
| --- | --- | --- |
| **Myh1** | AGCTGATGACCAACTTGCGC | CCCTGGAGACTTTGTCTCATTAGG |
| **Atrogin-1** | TAGGCGACCTCAGCAGTTAC | TGCAATATCCATGGCGCTCT |
| **β-actin** | AGGCACCAGGGCGTGAT | GCCCACATAGGAATCCTTCTGAC |
| **Myogenesis array kit** | AccuTarget™ qPCR Screening Kit (Bioneer SM-0148) | |

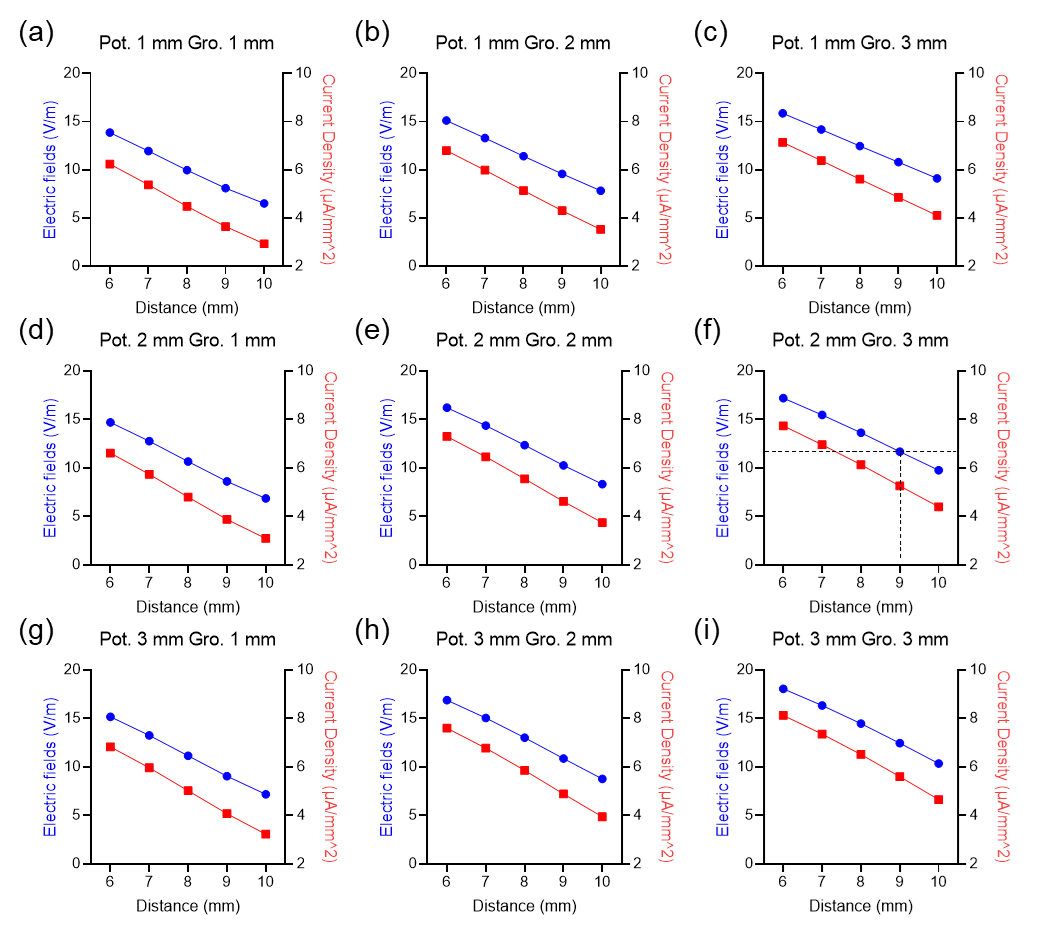

Fig. S1. Multiphysics simulation for translating in vitro to in vivo. (a-c) Simulation results based on the distance between the ground electrode and the potential electrode with the potential electrode width fixed at 1 mm. (d-f). Simulation results based on the distance between the ground electrode and the potential electrode with the potential electrode width fixed at 2 mm. (g-i) Simulation results based on the distance between the ground electrode and the potential electrode with the potential electrode width fixed at 3 mm.

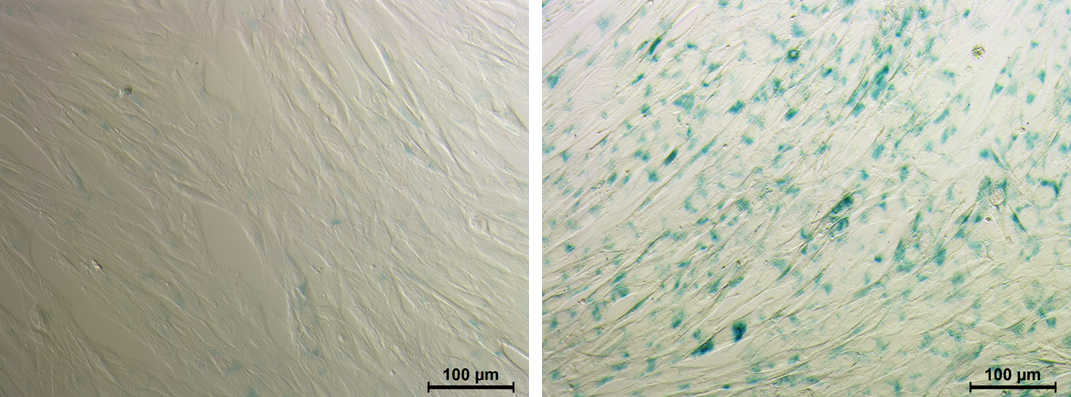

Fig. S2. SA-β-Galactosidase assay. (a) Young human skeletal muscle-derived cells are not stained by SA-β-Gal. (b) Representative image of SA-β-Gal positive senescent aged human skeletal muscle-derived cells (Dark blue color)
